# HSPB8 regulates CTP synthase filaments to couple nucleotide metabolism and autophagy in tumors

**DOI:** 10.64898/2026.02.10.704981

**Authors:** Chun-Yen Wang, Wei-Cheng Lin, Kuang-Jing Huang, Archan Chakraborty, Yu-Tsun Lin, Ya-Ju Hsieh, Kun-Yi Chien, Po-Yuan Ke, Wei-Han Huang, Chih-Yun Cheng, Ian Yi-Feng Chang, Hsiang-Yu Tang, Cheng-Hung Yang, Mei-Ling Cheng, Ying-Chih Chang, Chau-Ting Yeh, Chien-Kuo Lee, Jau-Song Yu, Yu-Sun Chang, Jörg Großhans, Li-Mei Pai

## Abstract

Metabolic adaptation and proteostasis are essential for tumor survival under nutrient stress, yet how these processes are mechanistically integrated remains unclear. Here, we identify CTP synthase (CTPS) filament dynamics as a regulatory nexus linking nucleotide metabolism to autophagic flux. Under glutamine deprivation—a hallmark of poorly vascularized solid tumors—CTPS undergoes polymerization into filamentous assemblies that exhibit reduced enzymatic activity. We demonstrate that filament formation is driven by intracellular asparagine availability and stabilized by the accumulation of misfolded proteins. Using APEX2-based proximity labeling and proteomics, we identify the small heat shock protein HSPB8 as a filament-associated regulator. HSPB8, acting within the chaperone-assisted selective autophagy (CASA) pathway, antagonizes CTPS polymerization by promoting clearance of misfolded proteins, thereby restoring soluble, catalytically active CTPS. Filament disassembly increases CTP production and enhances synthesis of autophagy-related phospholipids, including phosphatidylinositol and phosphatidylethanolamine, resulting in accelerated autophagic flux. Cells expressing filament-deficient CTPS mutants display elevated autophagosome formation and increased LC3-II accumulation upon lysosomal blockade, confirming enhanced flux. In vivo, disruption of CTPS filament assembly impairs xenograft tumor growth and is associated with excessive autophagy. Analysis of TCGA datasets further reveals that high CTPS and low HSPB8 expression correlate with poor patient survival across multiple cancers. Collectively, our findings establish CTPS filament dynamics as a proteostasis-sensitive metabolic switch that coordinates nucleotide biosynthesis with autophagy, revealing a previously unrecognized vulnerability in tumors adapting to nutrient limitation.

## Introduction

Adapting to chronic nutrient scarcity is a fundamental challenge for malignant progression. In solid tumors, poor vascularization leads to profound glutamine deprivation, forcing cancer cells to develop sophisticated survival strategies ^1^. Among these adaptive responses, cells increasingly rely on asparagine (Asn) to maintain proliferation under glutamine-depleted conditions ^2, 3, 4^. Consequently, asparaginase treatment is a key therapeutic strategy ^5, 6^. However, its efficacy is often limited by adaptive cellular responses, including the induction of autophagy ^7, 8^.

Emerging evidence suggests that cellular metabolism is regulated through the dynamic compartmentalization of enzymes into membrane-less organelles. Cytidine triphosphate synthase (CTPS), the rate-limiting enzyme in de novo pyrimidine synthesis, is a prominent example of this phenomenon. CTPS catalyzes the ATP-dependent conversion of UTP to CTP using glutamine as a nitrogen donor, and its expression and enzymatic activity have been established as drivers of malignant progression ^9^. Under glutamine-deprived conditions, mammalian CTPS undergoes a phase transition to form higher-order polymers and filament bundles, facilitated by specific residues in the linker (R294) and glutaminase (H355) domains ^10, 11^. Furthermore, we have previously shown that the histidine can trigger these filamentous structures to assemble along with cytokeratin networks and allow rapid cell growth after stress relief ^12, 13^. While these filaments are known to respond to nutrient cues, the molecular machinery that coordinates their assembly with cellular survival programs remains poorly defined.

The maintenance of proteostasis is primarily overseen by molecular chaperones, among which the small heat shock protein B8 (HSPB8) plays a specialized role. Unlike general chaperones, HSPB8 specifically recognizes unfolded peptides and defective ribosomal products (DRiPs), targeting them for degradation via the chaperone-assisted selective autophagy (CASA) complex ^14, 15, 16^. By forming a functional unit with the co-chaperone BAG3 and the scaffold protein HSP70, HSPB8 has been shown to actively promote autophagic flux to clear proteotoxic aggregates ^17, 18, 19^. While HSPB8 is well-characterized in the context of neurodegenerative protein aggregation, its involvement in regulating the structural dynamics of metabolic enzymes has not been explored. Given that metabolic stress often coincides with proteostatic imbalance, HSPB8 emerges as a potential link between the cellular protein quality control machinery and the adaptive compartmentalization of enzymes like CTPS.

In this study, we identify the availability of asparagine as a key environmental factor governing CTPS filament dynamics. Through proteomic screening, we discovered that HSPB8 serves as a critical metabolic rheostat that antagonizes CTPS polymerization. We demonstrate that filament assembly is driven by the accumulation of unfolded peptides, which act as structural scaffolds for filament bundling. Our results reveal that the HSPB8-mediated disassembly of CTPS filaments is a requisite step for activating autophagic flux under nutrient stress. By releasing soluble, active CTPS, cells increase CTP production to fuel the phospholipid biosynthesis required for autophagosome membrane expansion. Crucially, we show that disrupting this balance to reduce CTPS filament formation leads to significantly impaired tumor growth, identifying the HSPB8-CTPS-autophagy axis as a vital survival response. These findings define a therapeutic vulnerability in cancers with high CTPS expression and provide a new perspective on how cells coordinate their physical and chemical states to survive starvation.

## Results

### CTPS filament assembly is essential for xenograft tumor progression

Given that CTPS filaments have been identified in clinical tumor specimens ^20^, we hypothesized that these macromolecular structures serve a functional role in oncogenesis. To investigate whether CTPS polymerization regulates tumor growth *in vivo*, we performed xenograft experiments using HCT116 cells expressing either wild-type (WT) CTPS or the filament-deficient H355A mutant (**Figure 1A-B**). The filament-deficient H355A mutant cells exhibited a significant reduction in tumor volume compared to WT controls (**Figure 1C-E**). Notably, these phenotypic differences occurred in the absence of systemic toxicity, as indicated by stable murine body weights across cohorts (**Figure 1F**).

To further elucidate the mechanisms driving this growth disparity, we characterized the proliferative capacity of these cells under varying environmental conditions. Under glutamine deprivation in three-dimensional (3D) culture (Gln(-)S(+)), HCT116 cells showed negligible growth. However, in nutrient-replete conditions (Gln(+)S(+)), the H355A mutant displayed significantly reduced filaments and impaired proliferation specifically in suspension sphere assays (**Figure S1B-C**). This contrast is striking given that the mutant and WT cells exhibited indistinguishable growth rates in standard two-dimensional (2D) monolayer cultures (**Figure S1A**). Collectively, these data suggest that CTPS filament assembly is not merely a cellular feature but a critical regulator of 3D proliferative capacity and *in vivo* tumor formation.

### Asparaginase inhibits CTP synthase polymerization by depleting intracellular asparagine pools

Asparagine (Asn) is a known survival factor during glutamine deprivation ^3^. Consistent with this role, we observed that CTP synthase (CTPS) filament formation under glutamine-deprived conditions was inhibited by asparaginase (ASNase) treatment (**Figure 2, A to F**) without altering total CTPS protein levels (**Figure 2G**). Conversely, exogenous Asn supplementation promoted CTPS polymerization (**Figure 2, C to F**). To further define the specificity of this effect, we screened the remaining non-essential amino acids—alanine, asparagine, aspartate, and proline—for their capacity to induce filamentation. Our results confirm that Asn induces CTPS filaments, comparable to the histidine positive control (**Figure S2A**). Supporting this, the combination of Asn and ASNase treatment nearly abolished CTPS filamentation in EBSS, a conditional medium without any amino acid supplementation (**Figure S2B**).

Given that Asn is absent from DMEM, we hypothesized that serum serves as the primary source of Asn for filament assembly. Quantitative metabolic analysis revealed that Asn levels increase during glutamine deprivation relative to standard DMEM but are depleted upon serum withdrawal or ASNase treatment (**Figure S2C**). Although extracellular nutrients can be scavenged via macropinocytosis to support proliferation under nutrient stress ^21^, Asn can also be synthesized de novo by asparagine synthetase (ASNS). However, ASNS knockdown did not affect CTPS filamentation (**Figure S2, D and E**), likely due to substrate limitations of glutamine required for ASNS enzymatic activity ^22^. Collectively, these data demonstrate that intracellular Asn levels regulate CTPS filament dynamics, suggesting a metabolic adaptation where cells modulate amino acid homeostasis to confer nutrition stress during glutamine deprivation.

### Identification of CTPS-associated proteins via APEX2 proximity labeling

To elucidate the mechanisms governing CTP synthase (CTPS) filament dynamics, we employed APEX2-based proximity labeling to identify protein interactors in situ ^23^. Although CTPS filaments assemble during nutrient starvation, their structural integrity is highly sensitive to cell lysis, which typically precludes traditional biochemical purification. We therefore generated HEp-2 cell lines stably expressing a CTPS-APEX2 fusion protein to biotinylate proximal proteins in living cells ^13^. To ensure high specificity for filament-associated partners, we utilized two negative controls representing a non-filamentous state: glutamine-induced disassembly and a polymerization-deficient R294D mutant ^12, 24^ (**Figure 3A**). Western blot analysis confirmed that both endogenous CTPS and exogenous CTPS-APEX2 levels were markedly elevated under glutamine deprivation, consistent with reports that filamentation protects CTPS from proteasomal degradation ^12^. Furthermore, the enrichment of the known filament-associated protein IMPDH2 ^25^ in the CTPS-APEX2 pull-down validated the spatial accuracy and efficacy of our proximity labeling system (**Figure 3B**).

### Discovery of HSPB8 as a CTPS filament-associated protein regulated by asparaginase

To define the CTPS filament proteome with high stringency, we integrated data from two independent proteomics experiments and applied a fold-change cutoff based on IMPDH2 enrichment (**Figure S3, A to C**). This filtering strategy yielded 184 overlapping proteins significantly enriched in pathways involving de novo purine biosynthesis and protein ubiquitination (**Figure 3C**). IMPDH2 and PFAS in the purine de novo biosynthesis category have been demonstrated to form distinct compartments, filament, and purinosome, respectively ^26, 27^. Notably, the identification of multiple heat shock proteins (e.g., HSPA4, HSPB8, HSPH1) and deubiquitinating enzymes in the protein ubiquitination category suggests that protein quality control (PQC) machinery localizes proximally to CTPS filaments during metabolic stress. Given that asparagine (Asn) is a key effector of ATF4-mediated amino acid homeostasis, we hypothesized that Asn availability modulates this PQC-filament association ^4,28^. Quantitative proteomics and subsequent western blotting validation revealed that the small heat shock protein HSPB8 is downregulated during glutamine deprivation but specifically upregulated upon ASNase treatment (**Figure 3, D and E**). We further confirmed the physical proximity between CTPS and HSPB8 through a proximity ligation assay (PLA) (**Figure S3D**), establishing HSPB8 as a putative regulator of CTPS filament dynamics under glutamine-deprived stress.

### HSPB8 regulates the compartmentalization of CTPS filaments

HSPB8, a member of the small heat shock protein (sHSP) family, acts in concert with BAG3—and often within the chaperone-assisted selective autophagy (CASA) complex alongside HSP70 and E3 ligases—to facilitate the clearance of misfolded or aggregation-prone protein. Given its established role in maintaining proteostasis, we hypothesized that the ASNase-induced upregulation of HSPB8 promotes the disassembly of CTPS filaments. To test this, we performed loss- and gain-of-function assays to evaluate the requirement of HSPB8 for CTPS filament dynamics. Under glutamine-deprived conditions, HSPB8 knockdown significantly enhanced CTPS filament assembly compared to control groups (**Figure 4, A and B**). Conversely, we observed that the restoration of glutamine, which typically triggers filament disassembly, was less effective in the absence of HSPB8, resulting in persistent filament structures (**Figure 4, A and B**).

In contrast to the knockdown phenotypes, overexpression of HSPB8 markedly reduced the prevalence of CTPS filaments assembly, suggesting that HSPB8 facilitates the release of CTPS from filamentous compartments into the cytosol (**Figure 4, C and D**). Crucially, we found that HSPB8 depletion was sufficient to restore CTPS filament assembly even in the presence of ASNase treatment, which normally prevents filament assembly (**Figure S5A**). Collectively, these results demonstrate that ASNase disassembles CTPS filaments by upregulating HSPB8, identifying this chaperone as a key molecular rheostat for CTPS assembly during glutamine-deprived stress.

### Clinical significance of the HSPB8-CTPS axis in cancer prognosis

To evaluate the clinical relevance of the HSPB8-CTPS regulatory partnership, we first analyzed oral squamous cell carcinoma (OSCC) datasets (*n*=83) from the Chang Gung Memorial Hospital (CGMH) and the Molecular Medicine Research Center (MMRC) ^29^. In tumor tissues, *CTPS* mRNA levels were significantly elevated, whereas *HSPB8* mRNA levels were reduced compared to adjacent normal tissues (**Figure 4E**). Kaplan-Meier survival analysis of the CGMH-OSCC cohort indicated that patients with a low-*HSPB8*/high-*CTPS* expression profile exhibited a trend toward poorer overall survival (OS, *p*=0.68) and progression-free survival (PFS, *p*=0.25) (**Figure 4F**), though these results did not reach statistical significance likely due to the limited sample size.

To validate these findings in larger cohorts, we expanded our analysis to The Cancer Genome Atlas (TCGA) database. Across multiple malignancies—including BRCA, COAD, HNSCC, LUAD, LUSC, READ, STAD, and THCA—*CTPS* expression was significantly upregulated while *HSPB8* was markedly downregulated compared to normal controls (**Figure S4A and B).**

We further investigated the prognostic impact of this inverse correlation. Within the TCGA-HNSC cohort, among tumors with high *CTPS* expression, patients with low *HSPB8* levels showed a poorer prognosis compared to those with high *HSPB8* levels (*p*=0.054; **Figure 4G**). Strikingly, the combination of high *CTPS* and low *HSPB8* expression was a significant predictor of diminished survival in adrenocortical carcinoma (ACC), pancreatic adenocarcinoma (PAAD), sarcoma (SARC), and pheochromocytoma and paraganglioma (PCPG) (**Figure 4G and S4C**). Collectively, these clinical data suggest that an environment characterized by low *HSPB8*—which we have demonstrated promotes CTPS filament assembly—correlates with tumor progression and serves as a negative prognostic signature across several human cancers.

### Glutamine deprivation-induced misfolded protein accumulation drive CTPS filamentation

Since HSPB8 targets misfolded proteins for autophagic clearance and our proteomic data identified protein quality control (PQC) machinery in the CTPS interactome (**Figure 3C**), we hypothesized that the accumulation of unfolded proteins modulates CTPS filament dynamics. Proximity ligation assays (PLA) confirmed that the CTPS-HSPB8 association is spatially restricted to CTPS filaments (**Figure S5A**). Furthermore, we observed an increased association between HSPB8 and its co-chaperone BAG3 under glutamine deprivation (**Figure S5B**), suggesting an active role for the HSPB8-BAG3 complex in managing protein misfolding during metabolic stress.

To evaluate the direct impact of proteotoxic stress on CTPS polymerization, we treated cells with puromycin to induce the premature termination of translation and the subsequent generation of defective ribosomal products (DRiPs) ^30^. Under glutamine-deprived conditions, puromycin treatment resulted in the formation of significantly thicker CTPS filaments compared to untreated controls (**Figure 5, A and B**). The proximity biotin-labeling with streptavidin pull-down experiment revealed that puromycylated polypeptides are enriched in the immediate vicinity of CTPS filaments (**Figure 5C**), implying that local proteotoxic stress exerts a stabilizing effect on filament dynamics.

Consistent with its role in protein clearance, the knockdown of HSPB8 led to a marked accumulation of puromycylated polypeptides (**Figure 5D and Figure S5C**). Conversely, the intensity of these misfolded protein signals significantly declined upon ASNase treatment (**Figure 5E**). Taken together, these findings demonstrate that HSPB8 regulates CTPS filament dynamics by modulating the local concentration of unfolded proteins to create an environment for CTPS assembly, positioning the filament as a metabolic structure sensitive to cellular proteostasis.

### CTPS filament disassembly promotes autophagic flux under glutamine-deprived stress

Autophagy is a critical survival mechanism during asparaginase-induced cytotoxicity. Given that CTP is a requisite substrate for the de novo synthesis of choline phospholipids—catalyzed by PCYT1A (phosphate cytidylyltransferase 1, choline, alpha) to replenish autophagosome membranes ^31^—we investigated whether the disassembly of CTPS filaments enhances autophagic capacity. We previously established that non-filamentous CTPS possesses higher enzymatic activity than its polymerized counterpart. Consistent with this, our in vivo enzymatic assays, measuring the ^13^C^15^N-CTP to ^13^C^15^N-UTP ratio, demonstrated that filament-null mutants (R294D and H355A) exhibit significantly higher catalytic activity compared to wild-type CTPS (**Figure S6A**).

To determine if increased CTPS activity promotes autophagy, we utilized mRFP-GFP-LC3 reporter cell lines to monitor autophagic flux ^32^. Under glutamine deprivation, CTPS^R294D^ or CTPS^H355A^ cells displayed a higher density of mRFP-positive/GFP-negative puncta per cell than WT CTPS cells, indicating an acceleration of autolysosome formation (**Figure 6, A and B; Figure S6, B and C**). Transmission electron microscopy (TEM) further confirmed a marked accumulation of both early (AVi) and late (AVd) autophagic vacuoles in filament-null mutants, whereas wild-type cells exhibited significantly fewer structures (**Figure 6, C and D; Figure S6, D and E**). To distinguish between increased induction and blocked degradation, we treated cells with Bafilomycin A1 (BafA1) to inhibit lysosomal fusion. Under BafA1 treatment, filament-null cells accumulated significantly higher levels of LC3-II compared to wild-type controls, confirming that CTPS disassembly actively enhances autophagic flux (**Figure 6, E and F**). Collectively, these findings suggest that the metabolic regulation of CTPS filament dynamics modulates autophagic activity.

### HSPB8 promotes autophagy by antagonizing CTPS filament assembly

Building on findings identifying the chaperone HSPB8 as a negative regulator of CTP synthase (CTPS) polymerization, we investigated the role of CTPS filament dynamics within the HSPB8-autophagy axis during glutamine-deprived nutrient stress. HSPB8, a member of the small heat shock protein family, specifically binds unfolded peptides and has been previously implicated in the induction of autophagy ^33^. To determine if CTPS filament disassembly is a requisite step in this process, we evaluated autophagic flux under glutamine-and serum-deprived conditions (Gln(-)S(-))In cells expressing wild-type CTPS (WT CTPS), overexpression of HSPB8 significantly increased the number of autolysosomes (mRFP-positive/GFP-negative) and total autophagic vesicles compared to vehicle-treated controls (**Figure 7, A, C, and E**). In contrast, HSPB8 overexpression failed to further enhance autophagic vesicle accumulation in filament-deficient CTPS^H355A^ cells (**Figure 7, B, D, and E**). These results suggest that the pro-autophagic effect of HSPB8 under glutamine deprivation is mediated through the inhibition of CTPS filament assembly, identifying the transition from polymerized to dimeric or tetrameric CTPS as a critical metabolic switch for autophagic activation.

### Reduced volume of CTPS^H355A^ tumors correlates with excessive autophagy activity

Given that CTPS^H355A^ cells exhibit both impaired filament assembly and elevated autophagy in vitro, we sought to determine if this relationship holds true in a physiological tumor model. We analyzed the widely-used autophagy marker LC3B in WT CTPS – and CTPS^H355A^-derived tumors (from Figure 1) to understand how defective filament assembly interferes with tumor progression (**Figure S7A**). Consistently, the LC3B-II/LC3B-I ratio, a hallmark of autophagic flux, was elevated in CTPS^H355A^ tumors (**Figure S7B**). These findings suggest that the anti-tumor effect observed when CTPS filament assembly is disrupted is associated with an induction of excessive autophagic activity.

### Phospholipid biosynthesis is upregulated in filament-deficient CTPS mutants

Because CTPS facilitates phospholipid biosynthesis through CTP production, we hypothesized that filament-deficient mutants exhibit elevated autophagy due to an increased supply of membrane building blocks. Given that lipids and lipid-associated proteins are indispensable for autophagosome formation and maturation, we performed liquid chromatography-mass spectrometry (LC-MS) to compare the lipidomic profiles of WT CTPS and CTPS^R294D^ HEp-2 cells. We identified 75 annotated lipids that met the significance criteria of P < 0.05, fold change > 1.3, and a Variable Importance in Projection (VIP) score > 1 (**Figure 8A and Figure S8, A to C**). A heatmap of 43 high-confidence lipids which was verified by MS/MS identification revealed a consistent elevation in CTPS^R294D^ cells, correlating with the increased enzymatic activity of the non-filamentous mutant (**Figure 8B**). LIPEA pathway analysis indicated a significant enrichment of autophagy-related lipids, specifically phosphatidylethanolamines (PE) and phosphatidylinositols (PI), which were significantly higher in CTPS^R294D^ cells (**Figure 8, C and D**). Since PI and PE are essential for autophagosome membrane initiation and expansion, these findings suggest that the metabolic output of disassembled CTPS directly supports the lipid requirements for enhanced autophagic flux.

## Discussion

Metabolic enzymes increasingly are recognized for their ability to undergo higher-order compartmentalization in response to physiological stress, yet the functional consequences of these transitions remain poorly understood. Our findings demonstrate that the polymerization of CTP synthase (CTPS) into filamentous structures is a critical regulatory event that dictates tumor growth and metabolic adaptation. By integrating biochemical analysis, 3D cell culture, and clinical datasets, we show that CTPS filaments serve as a metabolic switch that balances enzyme activity with proteostatic demands (**Figure 1 and Figure S1**). Specifically, under glutamine deprivation—a hallmark of the nutrient-poor core of solid tumors ^1^—the assembly of CTPS filaments suppresses enzymatic activity, thereby modulating phospholipid availability and autophagic flux to influence tumor progression.

A key revelation of this study is the identification of the HSPB8-CTPS filament axis as a novel branch of the cellular protein quality control (PQC) system. Building on our previous evidence that ubiquitination regulates CTPS polymerization ^34, 35^, we identified a suite of PQC-related proteins associated with the filament interactome (**Figure 3**). The chaperone HSPB8, acting within the HSPB8-BAG3 complex, emerged as a master regulator of these dynamics. Our data suggest that misfolded proteins, or defective ribosomal products (DRiPs), localize to and stabilize CTPS filaments. By modulating the clearance of these proteotoxic species, HSPB8 dictates the architectural state of CTPS. The knockdown of HSPB8 promotes filament bundling, while its overexpression triggers disassembly (**Figure 4 A to F**). This mirrors the role of the HSPB8-BAG3-HSP70 complex in regulating the phase transition of stress granules, suggesting a universal role for HSPB8 in maintaining the fluidity and dynamics of biomolecular condensates and membrane-less organelles.

The biological significance of this regulation is underscored by our analysis of CGMH-OSCC and TCGA datasets, where we observed a pervasive inverse relationship between *CTPS* and *HSPB8* expression across 75% of cancer types (**Figure 4 E and S4, A to B**). The “high-*CTPS*/low-*HSPB8*” signature—which our model predicts would favor stable filament formation—correlates significantly with reduced patient survival (**Figure 4 F to G and S4 C**). This suggests that the stabilization of CTPS filaments may provide a selective advantage in aggressive malignancies, likely by sequestering enzymatic activity until metabolic conditions require its mobilization. Consequently, the molecular machinery governing CTPS assembly represents a promising, yet unexplored, therapeutic vulnerability in cancer.

Mechanistically, we establish that the transition from filamentous to soluble CTPS is a prerequisite for autophagic activation (**Figure 6 and Figure S6**). While it is well known that autophagy is essential for surviving starvation, we found that the pro-autophagic effects of HSPB8 are strictly dependent on the filamentous state of CTPS (**Figure 7**). The disassembly of these structures facilitates CTP production, providing the necessary energy and substrates for phosphatidylcholine biosynthesis, which is required for autophagosome membrane expansion. Our lipidomic profiles confirm that autophagy-essential lipids, such as PI and PE, are upregulated upon filament disassembly (**Figure 8**). In the context of the tumor microenvironment, the mobilization of CTPS and the resulting surge in autophagy may interfere with tumor growth (**Figure S7**), as excessive autophagic flux can trigger apoptosis or metabolic exhaustion ^36, 37^.

Finally, our study identifies the structural dynamics of CTPS as a direct downstream target of the cellular PQC response. We propose that the accumulation of unfolded peptides serves as a molecular scaffold that promotes CTPS filament bundling. This process is highly sensitive to the cellular methylation status ^12^, as evidenced by the disassembly of CTPS filaments upon treatment with the methylation inhibitor Adox (**Figure S2F**). Under metabolic stress and asparaginase treatment, the upregulation of HSPB8 facilitates the clearance of these filament-associated misfolded proteins (**Figure 2 and Figure 3E**). This HSPB8-mediated transition, potentially modulated by methylation-dependent signaling ^38^, ensures that cells can reallocate metabolic resources from a sequestered state into active CTP synthesis to meet the demands of autophagic flux. Collectively, these findings position the CTPS filament as a structural bridge between protein quality control and metabolic homeostasis, revealing that targeting the unfolded protein-HSPB8 axis could disrupt the safeguards enabling cancer cell survival (**Figure 8E**).

## Material and Methods

### Cells

Human HEp-2 and HeLa cells were cultured in Dulbecco’s Modified Eagle’s Medium (Gibco) supplemented with 10% fetal bovine serum. Human HCT116 cells were culture in McCoy’s 5A Medium (Sigma). All of cells were maintained at 37°C under 5% CO^2^, and passage was done every 3∼4 days. Cells were wash with 1X PBS twice before any medium change.

### 3D spheroid culture

HCT116 cells were seeded in the R^3^CE platform (Acrocyte Therapeutics Inc., New Taipei City, Taiwan) and incubated in McCoy’s 5A medium (with 10% FBS), and appropriate amount of fresh medium is replenished every 3-4 days. Spheroid images were captured by Nikon Eclipse-TS100 microscope with LeadView-2000AIO camera and the spheroid sizes were quantified by imageJ. The spheroids at day 21 were fixed by 4% formaldehyde for 1 h 30 mins and immunostained with CTPS, followed with image capturing by Leica Stellaris SP8 confocal microscope.

### Xenograft mouse

Fresh HCT116 WT-CTPS and H355A-CTPS are suspended in McCoy’s 5A (without FBS) with Matrigel (M5A: Matrigel=1:1) and subcutaneously injected into right dorsal flank of NUDE mice (BALB/cAnN.Cg-Foxn1nu/CrlNarl). From tumor start to be seen, tumor volume is calculated by length and width (tumor volume V = ½ (Length × Width2)) twice a week. Until 28 days after injection, mice are sacrificed and tumors are collected for further experiments.

### Plasmid construction

To conduct the cells which stably express CTPS1-APEX2 gene, the CTPS1-APEX2 was amplified from p3xFlag-CTPS1-APEX2-CMV26 vector (*13*) and inserted into MIGR1 plasmid with XhoI sites. The mutants (CTPS^R294D^ and CTPS^H355A^) were generated by primers harboring the desired mutations through site-directed mutagenesis. To generate the cells which stably express CTPS1-AcGFP, the CTPA1-AcGFP gene was amplified from p3xFlag-CTPS1-GFP-CMV26 vector (*13*) and inserted into pLAS2w.Ppuro vector with NheI sites. Amplification of PCR products by KOD DNA Polymerase (Toyobo). Plasmids were transfected into HEp-2 or 293T cells using Lipofectamine 2000 (Invitrogen) following the manufacturer’s instructions.

### Immunofluorescent staining

5 x 10^4^ human HEp-2 or HeLa cells were seeded onto coverslips in 24-well plates and incubated overnight. For HCT116 cells, cells were seeded and incubated for 48 h. Culture medium was replaced with conditional medium (glutamine negative DMEM and serum/glutamine negative DMEM for 24 h, EBSS and EBSS + amino acids for 4 h). Cells were washed with PBS (containing Mg^++^ and Ca^++^) once and fixed in 4% formaldehyde for 10 min and 100% methanol (-20) for 2 min, followed by permeabilization with blocking buffer (2% BSA and 0.2% Triton X-100 in PBS) for 20 min. Cells were incubated with primary antibody overnight at 4 and secondary antibody for 1 h. Coverslips were mounted in mounting medium and samples examined by confocal microscopy (Zeiss LSM 510 Meta confocal microscope).

### Quantification of CTPS filament formation

The quantification of CTPS filament formation was followed with our previous publication in 2018^13^ and 2022^40^. For the percentage of filament bearing cells, images were captured from at least three random fields of each individual immunostaining coverslips, and the numbers of CTPS filament-bearing cells were normalized with the total cell numbers in each images to obtain the percentage value that represented the filament forming abilities in every group of experiment. Three biological repeats of the percentage values were summarized in the bar charts next to the representing immunostaining images. For intensity of CTPS filament (in figure 4 B), the intensities of green filaments (CTPS filaments) were measured by Fiji (ImageJ). Every dot represents the intensity of one CTPS filament, and the intensity values from multiple-field images of three biological repeats were summarized in the scatter plot which represented the size distribution of CTPS filament.

### Western blotting

Human HEp-2 cells (3 x 10^5^) were seeded on 3.5-cm culture dishes and incubated overnight. Culture medium was replaced with conditional medium for the indicated times. Cell lysates or small tumor dissection were extracted with RIPA buffer (50mM Tris-HCl [pH 7.2], 150mM NaCl, 2mM EDTA, 1% NP40, 0.1% SDS, 1% sodium deoxycholate) and 1X protease inhibitor cocktail (Complete Mini, Roche). Equal amounts of lysates (20μg) were loaded per well, subjected to SDS-PAGE, and transferred to PVDF membranes. Membranes were blocked with 5% nonfat milk or 5% BSA in TBS-T (0.1% Tween 20 in Tris-buffered saline) for 1 h and incubated with primary antibodies diluted 1:1000 in blocking buffer at 4, followed by incubation of secondary antibodies for 1 h. Membranes were developed by ECL (PerkinElmer), signals of bands were detected on a transilluminator (UVP) and quantified by densitometry.

### shRNA gene knockdown

The lentivirus-based shRNA knockdown system was provided by the National RNAi Core Facility, Academia Sinica. The cells (3 x 10^5^) were infected with 20μl recombinant viral fluid in the presence of 8 mg/ml polybrene (Sigma-Aldrich) for 24 h. Infected cells were grown in fresh DMEM with 2 μg/ml puromycin (Invivogen) every 48 h for one week. Puromycin resistant cells were collected for experiments. Efficiency of knockdown on specific gene was validated by Western blotting.

### Gene expression

6 x 10^5^ HEp-2 cells were seeded on 6-cm culture dishes and incubated overnight. Culture medium was replaced with conditional medium for the indicated times. Total RNA was isolated from cells by TRIzol Reagent (Thermo Fisher Scientific), followed by reverse transcription with RevertAid First Strand cDNA Synthesis Kit (Thermo Fisher Scientific). Real-time PCR amplification was performed using specific primers and Power SYBR Green PCR Master Mix (Thermo Fisher Scientific) in a StepOnePlus instrument (Thermo Fisher Scientific). Relative expression of target genes was obtained by normalization of control group. Statistical analyses were performed by Student’s t test.

### Measurement of labeling UTP and CTP

Analysis was performed as previously described (*12*). 1.5 x 10^6^ cells were harvested by methanol. Supernatants were dried and re-suspended in 20 mM Dibutylamine and 20 mM Formic acid. Extracts of 10^4^ cells were analyzed by LC-MS (LTQ-orbitrap Elite, Thermo Fisher Scientific, Waltham, MA, USA) linked to a Waters ACQUITY UPLC system (Waters, Milford, MA, USA) connected with Hypresil Gold C18 (2.1 X 50 mm, 1.9 um particle size, 175 Å, Thermo Fisher Scientific). The flow rate is 60 μl/min and solvent A was 2 mM Dibutylamine and 2 mM Formic acid in water, and solvent B was Methanol with 20 mM Dibutylamine. The gradient elution conditions were: t = 0 min, 2% B; t = 1 min, 8% B; t = 11 min, 13% B; t = 14 min, 95% B; t = 16 min, 95% B; t = 16.5 min, 2% B; t = 20 min, 2% B. All MS spectra were obtained in the negative ion mode. The peak area of ^12^C-CTP/UTP and ^13^C-CTP/UTP from the raw LC-MS data were analyzed using Thermo Xcalibur Qual Browser 2.2 software (Thermo) (*39*).

### Quantification of hydrophilic metabolites with liquid chromatography coupled with tandem mass spectrometry

After metabolism quenching, cell suspension is vortexed, centrifuged at 12000 g for 15 min. The supernatant can be dried under nitrogen gas, dissolved in 100 μL water, and centrifuged at 12,000 g for 30 min to remove debris. The supernatant were analyzed using Waters ultra-high-performance liquid chromatography coupled with Waters Xevo TQXS MS (Waters Corp.). MS was operated in negative mode. The optimized parameters were as follows: capillary voltage at 1 kV; desolvation temperature at 500°C; source temperature at 150°C; and gas flow at 1000 L/h. The chromatographic separation was achieved on a BEH C18 (100 x 2.1 mm, particle size of 1.7 um; Waters Corp.) at 45°C with elute A (water with 10 mM tributylamine and 15 mM acetic acid) and eluent B (50% acetonitrile with 10 mM tributylamine and 15 mM acetic acid), and the flow rate was set at 0.3 mL/min. The gradient profile was as follows: iso-gradient 4% B, 6min; linear gradient 4-50% B, 0.1 min; 50%-60% B, 2.9 min; 60-100% B, 0.8 min, and keep 2.2 min. Chromatographic separation was performed on a C18 column (2.1 mm × 100 mm × 1.7 µm, Waters corp.). Mix QC sample (a mixture of all samples) was prepared for analyzed during the analytical runs after every 10^th^ sample.

### Proliferation assays

1 x 10^5^ HEp-2 cells were seeded in triplicate in 3.5-cm petri dishes overnight. For HCT116 cells, 1 x 10^5^ were seed for 48 h. Cells (cells numbers were labeled with dash lines) replaced with indicated conditional medium for 48 h. then cells were counted by trypan blue exclusion assay.

### PLA

Samples were prepared as previously described (*13*). A total of 5 × 10^4^ HEp-2 cells were seeded for experiments. After fixation (4% formaldehyde and 4% sucrose in 1× PBS) and permeabilization (100% ice-cold methanol), PLA (Duolink PLA, DUO92101 Sigma) was performed based on the manufacturer’s protocol.

### Transmission electron microscopy

Cells were seeded on the ACLAR^®^ 33C film (EMS) overnight and then treated with the desired condition. Films are fixed in 2.5% glutaraldehyde followed by post-fixation of 1% osmium tetroxide (EMS) solution for 1 hrs. After dehydration in graded series of ethanol, we inversed the film on the top of the resin block thus embedding the intact cell monolayer of adhesion cells. Monolayers were embedded in the Spurr’s resin (EMS) and then polymerized in an oven at 70°C for 8 h. Ultrathin sections (70 nm) were cut using Leica UC7 ultramicrotome, and collected onto a copper grid for TEM observation. Finally, the sections were post-stained by 4% uranyl acetate for 10 minutes, and rinsed several times with H2O followed by 4% Reynolds lead citrate for 10 minutes. Micrographs were obtained at 100 kV in a JEM-1230 transmission electron microscopy (JEOL) with a Gatan Model 832 digital camera. Morphology of autophagic vacuoles was considered from other studies (*40*).

### Trypsin digestion for 6-plex Tandem Mass Tag (TMT) labeling

For CTPS-APEX2 pull-down experiments, biotinylated proteins were pulled down using streptavidin-coated magnetic beads. The bound proteins were eluted using 80% trifluoroethanol (TFE)/0.1% trifluoroacetic acid (TFA) and dried in a speed vac. For whole cell lysates experiments, cultured cells were homogenized in RIPA buffer (50 mM Tris-HCl, 150 mM NaCl, 2 mM EDTA, 1% NP40, 0.1% SDS, 1% sodium deoxycholate, pH 7.2) containing 1X protease inhibitor cocktail on ice, and followed by sonication and centrifugation at 12,000 rpm for 30 min at 4°C. Supernatants were collected and stored in aliquots at -80°C after assessing protein concentration using a BCA assay kit (Pierce, Rockford, IL, USA). Dried pull-down products or cell lysates were dissolved in 250 mM triethylammonium bicarbonate (TEABC), reduced with 5 mM tris(2-carboxyethyl)phosphine (TCEP) at 60°C for 1 hr, alkylated with 10 mM methyl methanethiosulfonate (MMTS) for 30 min at RT and digested with sequencing grade modified porcine trypsin (20 μg/ml) (Promega, Madison, WI, U.S.A) overnight at 37°C. The tryptic digested peptides were vacuum dried, dissolved in 100 mM TEABC and subjected to TMT labeling according to the manufacturer’s instruction (Thermo-Fisher Scientific, Waltham, MA, U.S.A). Labeled peptides were mixed and further desalted for two-dimensional liquid chromatography-tandem mass spectrometry (2D-LC-MS/MS) analysis.

### 2D-LC-MS/MS analysis

The TMT6-labeled mixtures were separated and analyzed by 2D-LC-MS/MS using a strong cation exchange (SCX) and reverse-phase C18 (RP18) liquid chromatography system on a Dionex UltiMate 3000 nano LC system coupled to a LTQ-Orbitrap Elite mass spectrometer (Thermo-Fisher Scientific). The peptide mixture was reconstituted in HPLC buffer A (30% acetonitrile/0.1% formic acid) and loaded onto a homemade SCX column (Luna SCX, bead size, 5 μm; column dimensions, 180 × 0.5 mm) (Phenomenex Inc., Torrance, CA, USA) at flow rate of 5 μl/min for 30 minutes. The peptides were then fractionated into 48 fractions for the CTPS-APEX2 pull-down experiments, and 72 fractions for whole cell lysates using a continuous HPLC buffer B gradient (0-100% of 0.5 M ammonium chloride in the presence of 25% acetonitrile/0.1% formic acid). Each fraction was then mixed with a stream of 0.1% FA/H2O, and the peptides were trapped on a Zorbax C18 column (bead size, 5 μm; pore size, 30 nm; column dimensions, 5 × 0.3 mm) (Agilent Technologies Inc., Santa Clara, CA, USA) and separated with a 60-min acetonitrile gradient in the presence of 0.1% formic acid on a BEH C18 nanoACQUITY column (bead size, 1.7 μm; pore size, 12 nm; column dimensions, 100 × 0.1 mm) (Waters Corporation, Milford, MA, U.S.A).

MS/MS analysis was performed on an LTQ-Orbitrap Elite mass spectrometer (Thermo-Fisher Scientific). Full-scan MS spectra (m/z 400 - m/z 1600) were acquired in the orbitrap analyzer at a resolution of 60,000 at m/z 400. For CTPS-APEX2 pull-down experiments, the ten most intense precursor ions (intensity threshold of 10,000) were selected for fragmentation by higher energy collisional dissociation (HCD) with a normalized collision energy setting of 35% and an activation time of 0.1 ms. The dynamic exclusion function was set as: repeat count, 2; repeat duration, 30s; and exclusion duration, 60s. For whole cell lysates, the eight most intense precursor ions, above a threshold of 20,000, were selected for fragmentation by collision induced dissociation (CID), in addition to HCD, with a normalized collision energy setting of 35% and an activation time of 10 ms for CID and 0.1 ms for HCD. The dynamic exclusion function was set as: repeat count, 2; repeat duration, 30 sec.; and exclusion duration, 40 sec.

### Database search of 2D-LC-MS/MS

The MS and MS/MS data were analyzed and quantified using Proteome Discover (version1.4) (Thermo-Fisher Scientific) and processed with Mascot (version 2.5.1), matching against the Swiss-Prot database (download in January 2020; containing 20,368 human entries). Mass tolerance for parent and fragment ions was set to 10 ppm and 1 Da for CID and 10 ppm and 0.1 Da for HCD (0.05Da for CTPS-APEX2 pull-down experiments, respectively. Two missed cleavages were allowed for trypsin digestion; S-methylthio-cysteine was set as a fixed modification; oxidation of methionine, protein N-terminal acetylation, glutamine to pyroglutamic acid conversion at peptide N-terminus, and TMT labeling of lysine and N-termini of peptides were set as variable modifications. The criteria for peptide identification were as follows: peptide confidence, high; peptide length, 7–100; peptide maximum rank, 1; search engine rank, 1; minimal number of peptides for a protein group, 2; count only rank 1 peptide; count peptides only in top-scoring proteins; and FDR < 0.01.

### Hydrophobic metabolite profile

The cell samples were quenched with 1 mL of 80% methanol. The 300 µL sample was mixed with 30 µL methanol and 900 µL methyl tert-butyl ether (MEBE)and vortex for 1 min, and then stay at room temperature for 1 hour for protein precipitation. Sample were added into 156 µL water, vortex for 1 min, and stay at room temperature for 10 min. Samples were centrifuged with 12000 g for 30 min at 4°C, and 700 µL supernatant were transfer to a new microtube. The residual sample was extracted again with 700 µL MTBE, and then stay at room temperature for 1 hour. Sample were centrifuged with 12000 g for 30 min at 4°C, and 700 µl supernatant were transferred to previous tube and dried with nitrogen gas. Before lipidomics analysis, sample were redissolved in 1 mL isopropanol/acetonitrile/water (2:1:1, v/v/v) mixture, and were centrifuged 12000 g for 30 min at 4°C. The supernatant was dropped to lipidomic analysis with ultra-high-performance liquid chromatography coupled with Synapt G2-S (Waters Corp., Milford, MA, USA). MS was operated in both positive and negative mode. The optimized parameters in both modes were as follows: capillary voltage at 1.5 kV; desolvation temperature at 550°C; source temperature at 120°C; and gas flow at 900 L/h. The chromatographic separation was achieved on a BEH C18 column (100 x 2.1 mm, particle size of 1.7 µm; Waters Corp., Milford, MA, USA) at 60°C with elute A (acetonitrile/water (40:60, v/v) with 10mM ammonium formate) and eluent B (isopropanol/acetonitrile (90:10, v/v) with 10mM ammonium formate), and the flow rate was set at 0.45 mL/min. The gradient profile was as follows: linear gradient 40%-99% B, 10 min. The column was then re-equilibrated for 2 min for next analysis. Mix QC sample (a mixture of all samples) were prepared for analyzed during the analytical runs after every 10th sample.

### Lipid metabolite analysis

The methodological flow chart of lipid candidates narrowing down is summarized in S8A. Briefly, Original lipid targets were analyzed by VIP score, fold change, and abundance for the first cut-off. Then the 453 differently altered metabolites were preliminarily annotated by mass-to-charge ratio (m/z), partition coefficient (logP), and retention time screening (in HMDB), and they were secondly narrowed down by statistical significance (p-value < 0.05). 43 out of 75 narrowed metabolites were verified by MS/MS identification and was listed in Fig S8C. These verified metabolites (43 candidates) were then analyzed in MetaboAnalyst (https://www.metaboanalyst.ca/) and LIPEA pathway annotation tool (https://hyperlipea.org/home).

### TCGA data analysis

TCGA cancer cohorts were used for CTPS and HSPB8 expression levels screening between multiple cancer types. The Kaplan–Meier survival plots were analyzed with two subgroups categorized by gene expression: CTPS-high/HSPB8-high and CTPS-high/HSPB8-low in each type of cancer.

### Statistical Analysis

Experiment data are analyzed by Student’s t test or Kaplan-Meier analysis. Results includes at least 3 biological repeats show means and standard deviation (SD) values with P-values: *<0.05, **<0.01, ***<0.001, ****<0.0001. Other statistical details are included in figure legends.

## Use of Generative AI

Original writing was done by W.C.L., C.Y.W., A.C., and L.M.P.. Gemini (Google) and ChatGPT (OpenAI) were used to improve the quality of the writing for a specific section(s) of the manuscript.

## Acknowledgements

We thank Chang Gung University for the Metabolomics Core Laboratory for assistance with metabolomics analysis, Proteomics Core Laboratory for assistance with proteomic analysis, Microscopy Center for assistance with image capture, TEM and ultramicrotomy, Bioinformatics Core Laboratory for assistance with TCGA database analysis, respectively. We thank the National RNAi Core Facility at Academia Sinica for providing shRNA reagents and related services. We thank Dr. Hui-Kuan Lin, Anna C-C Jang and Tzyy-Choou Wu for their comments on the manuscript.

## Author Contributions

Conceptualization: W.C.L. and L.M.P. Methodology: K.Y.C., M.L.C., P.Y.K., and C.K.L.

Investigation: W.C.L., C.Y.W., K.J.H., A.C., W.H.H., C.Y.C., Y.T.L., Y.J.H., and H.Y.T.

Validation: A.C.

Visualization: W.C.L., C.Y.W., K.J.H., and A.C.

Data curation: W.C.L., C.Y.W., K.Y.C., Y.T.L., C.H.Y., and M.L.C.

Formal analysis: Y.S.C. and Y.F.C.

Funding acquisition: W.C.L. and L.M.P.

Project administration: L.M.P.

Supervision: Y.S.C. and J.S.Y.

Resources: P.Y.K., C.T.Y., Y.C.C., and C.K.L.

Writing – original draft: W.C.L., C.Y.W., A.C., and L.M.P.

Writing – review & editing: Y.S.C., C.K.L., and J.G.

## Data availability statement

Information and requests for resources and reagents should be directed to and will be fulfilled by the corresponding author, Dr. Li-Mei Pai. The mass spectrometry proteomics data of CTPS-APEX2 filament proximity labeling have been deposited to the ProteomeXchange Consortium via the PRIDE ^39^ partner repository with the dataset identifier PXD036474. The CGMH OSCC patient transcriptomics data were available in the previous publication ^29^.

## Abbreviation list

Adox: adenosine dialdehyde,
APEX2: enhanced ascorbate peroxidase 2
BAG3: BAG cochaperone 3,
CTP: cytidine triphosphate,
CTPS: cytidine triphosphate synthase
DMEM: Dulbecco’s modified eagle Medium,
H355A: histidine at the 355^th^ amino acid of CTPS was replaced with alanine
HSPB8: heat shock protein family B (small) member 8,
GFP: green fluorescent protein,
IMPDH2: inosine-5’-monophosphate dehydrogenase 2,
LC3 (MAP1LC3): microtubule-associated protein 1 light chain 3
M5A: McCoy’s 5A medium,
mRFP: monomeric red fluorescent protein,
R294D: arginine at the 294^th^ amino acid of CTPS was replaced with aspartic acid
UTP: uridine triphosphate.

## Figure Legends

**Figure 1. Disrupting CTPS filament assembly impairs HCT116 tumor growth in vivo**

**(A)** HCT116 cells stably expressing WT CTPS or CTPS^H355A^ were cultured in Gln-free DMEM for 24 h, and immunostained for CTPS (green); nuclei were counterstained with DAPI (blue). **(B)** Schematic of the xenograft mouse experiment. Thirty mice were used, with 15 injected with HCT116 WT CTPS cells and 15 injected with HCT116 CTPS^H355A^ cells. The experiment was terminated day 28 after injection. **(C)** Tumor volume measured by external caliper throughout the experimental period. **(D)** Representative images of tumors derived from WT CTPS or CTPS^H355A^ cells excised at day 28 after injection; tumor weights are quantified in **(E)**. **(F)** Mouse body weight recorded by scale over the course of the experiment.

**Figure 2. Asparagine availability is essential for CTPS filament assembly under glutamine deprivation**

**(A, B)** HCT116 cells were incubated in Gln-free DMEM or Gln-free DMEM supplemented with ASNase (2U/ml) for 24 h, followed by immunostaining for CTPS (green) and DAPI (blue). **(C, D)** HEp-2 cells were incubated in Gln-free DMEM supplemented with ASNase or Asn (0.34mM) for 24 h, followed by immunostaining for CTPS (green) and DAPI (blue). **(E, F)** HeLa cells were incubated with Gln-free DMEM supplemented with ASNase or Asn (0.34mM) for 24 h, followed by immunostaining against CTPS (green) and DAPI (blue). **(G)** HEp-2 cells were incubated in indicated conditions for 24 h, harvested and lysed for Western blotting analysis. Scale bars: 10 mm.

**Figure 3. Proximity labeling identifies the protein quality control factor HSPB8 as a condition-specific interactor of CTPS**

**(A)** Schematic of the CTPS-APEX2 proximity labeling strategy. **(B)** HEp-2 cells expressing WT CTPS-APEX2 were cultured in DMEM, Gln-free DMEM, or Gln-free DMEM supplemented with Gln for 24 h. HEp-2 cells expressing CTPS^R294D^-APEX2 were cultured in Gln-free DMEM for 24 h. Biotinylated proteins were affinity purified using streptavidin-conjugated magnetic beads and analyzed by Western blotting. **(C)** Venn diagram showing 184 proteins common to the comparisons of WT CTPS-APEX2 cells cultured in Gln-free DMEM versus Gln-free DMEM supplemented with Gln, and WT CTPS-APEX2 versus CTPS^R294D^-APEX2 cells cultured in Gln-free DMEM. Enriched pathways associated with CTPS-interacting proteins under Gln deprivation were identified by Ingenuity Pathway Analysis (IPA). **(D)** Heatmap showing proteins involved in the unbiquitination pathway in cells cultured in DMEM, Gln-free DMEM, or Gln-free DMEM supplemented with ASNase. **(E)** HSPB8 protein levels in cells cultured in DMEM, Gln-free DMEM, or Gln-free DMEM supplemented with ASNase.

**Figure 4. HSPB8 is required for the disassembly of CTPS filaments during nutrient stress**

**(A and B)** HCT116 cells expressing shLacZ or shHSPB8 were cultured in glutamine-free DMEM for 24 h, with glutamine added for the final 30 min where indicated. Cells were immunostained for CTPS (green) and DAPI (blue). CTPS filament intensity was quantified using ImageJ, with representative filament lengths indicated by pixel values (yellow). HSPB8 protein levels were assessed by Western blotting.

**(C and D)** HCT116 cells transfected with HA-mock or HA-HSPB8 were cultured in glutamine-free DMEM for 24 h and immunostained for CTPS (green) and DAPI (blue).

**(E and F)** Patient analysis of CGMH-OSCC dataset. Violin plot showed that the mRNA levels of CTPS and HSPB8 (**E**) and the Kaplan–Meier overall survival analysis comparing HSPB8 low and high expression in high CTPS expression cohorts (**F**).

**(G)** Kaplan–Meier overall survival analysis of TCGA datasets indicating that tumors with high CTPS and low HSPB8 expression are associated with significantly reduced survival, compared to that with high CTPS and high HSPB8 expression, in head and neck squamous cell carcinoma (HNSC), adrenocortical carcinoma (ACC), pancreatic adenocarcinoma (PAAD), and pheochromocytoma and paraganglioma (PCPG).

**Figure 5. HSPB8-mediated protein quality control is associated with CTPS filament formation**

**(A and B)** HCT116 cells were cultured in Gln-free DMEM for 24 h, with puromycin added during the final 3 h, followed by immunostaining for CTPS (green) and DAPI (blue). CTPS filament intensity was quantified using ImageJ.

**(C)** HCT116 cells expressing WT CTPS-APEX2 were cultured in Gln-free DMEM for 24 h, with puromycin (10 μg/ml) added during the final 3 h. Proximity labeling was performed, and biotinylated proteins were analyzed by Western blotting.

**(D)** shLacZ- and shHSPB8-expressing HEp-2 cells and **(E)** HEp-2 cells were cultured in Gln-free DMEM for 24 h, with puromycin (20 μg/ml) added during the final 10 min. Cells were harvested and analyzed by Western blotting for puromycin incorporation, HSPB8, and actin.

Scale bars, 10 μm.

**Figure 6. CTPS filament modulates autophagic flux under glutamine deprivation**

**(A and B)** HEp-2 cells stably expressing WT CTPS or CTPS^R294D^ and the tandem GFP–RFP–LC3 reporter were cultured in Gln-free DMEM for 24 h. GFP- and mRFP-positive puncta were quantified using ImageJ.

**(C and D)** Transmission electron microscopy images of WT CTPS and CTPS^R294D^ HEp-2 cells cultured in Gln-free DMEM for 24 h. Autophagic vesicles are indicated by yellow arrowheads.

**(E and F)** Quantification of LC3-II levels in WT CTPS and CTPS^R294D^ HEp-2 cells cultured in Gln-free DMEM for 24 h, with or without bafilomycin A1 (BafA1; 100 nM) added during the final 16 h.

**Figure 7. HSPB8 overexpression enhances autophagy inversely correlated with CTPS filament formation**

**(A–D)** Representative images of autophagic vesicles in HCT116 cells expressing the tandem RFP–GFP–LC3 reporter. Red-only puncta indicate autolysosomes, while yellow puncta (RFP/GFP colocalized) mark autophagic structures; nuclei were counterstained with DAPI (blue).

**(A′–D′)** Corresponding images showing CTPS filament formation by immunostaining for CTPS (green) with DAPI (blue).

**(E)** Quantification of autophagic puncta per cell, including red-only, yellow, and total puncta. Numbers below each condition (n) indicate the number of cells analyzed per group.

Scale bars, 10 μm.

**Figure 8. Metabolomic profiling of phospholipids in cells expressing WT CTPS versus filament-null CTPS mutant**

**(A)** Scheme of the experiment design. HEp-2 cells stably expressing WT CTPS and CTPS^H355A^ were cultured in Gln-free DMEM for 24 h, and subjected to lipid extraction, followed by LC/MS-MS analysis. Three independent experiments were performed and included in the analysis. **(B)** Heatmap showing the relative abundance of 43 identified metabolites across conditions. **(C)** Pathway annotation analysis of the 43 identified metabolites using LIPEA. **(D)** Phosphatidylserine (PS) and Phosphatidylethanolamine (PE) contribution to autophagy pathway. **(E)** Schematic illustrating the HSPB8–CTPS axis reveals a metabolic vulnerability in glutamine-stressed tumors. CTPS filament dynamics is suggested as a metabolic switch. Accumulation of unfolded peptides promotes CTPS filament bundling, a process modulated by cellular methylation. Under glutamine deprivation, reduced HSPB8 levels lead to the accumulation of misfolded peptides, thereby facilitating CTPS filament assembly. In contrast, increased HSPB8 expression, induced by asparaginase (ASNase), promotes CTPS filament disassembly and enhances CTPS enzymatic activity. Elevated CTPS activity supports CTP-dependent phospholipid synthesis, including phosphatidylinositol (PI) and phosphatidylethanolamine (PE), thereby driving autophagosome expansion and suppressing tumor growth. This regulatory axis reveals a metabolic vulnerability in tumors characterized by high CTPS and low HSPB8 expression.

Abbreviations: Me, methylation; ASNase, asparaginase.

## References

1. Pan M, Reid MA, Lowman XH, Kulkarni RP, Tran TQ, Liu X, et al. Regional glutamine deficiency in tumours promotes dedifferentiation through inhibition of histone demethylation. Nat Cell Biol 2016, 18(10): 1090–1101.

2. Krall AS, Xu S, Graeber TG, Braas D, Christofk HR. Asparagine promotes cancer cell proliferation through use as an amino acid exchange factor. Nat Commun 2016, 7: 11457.

3. Pavlova NN, Hui S, Ghergurovich JM, Fan J, Intlekofer AM, White RM, et al. As Extracellular Glutamine Levels Decline, Asparagine Becomes an Essential Amino Acid. Cell Metab 2018, 27(2): 428–438 e425.

4. Ye J, Kumanova M, Hart LS, Sloane K, Zhang H, De Panis DN, et al. The GCN2-ATF4 pathway is critical for tumour cell survival and proliferation in response to nutrient deprivation. EMBO J 2010, 29(12): 2082–2096.

5. Lorenzi PL, Claerhout S, Mills GB, Weinstein JN. A curated census of autophagy-modulating proteins and small molecules: candidate targets for cancer therapy. Autophagy 2014, 10(7): 1316–1326.

6. Willems L, Jacque N, Jacquel A, Neveux N, Maciel TT, Lambert M, et al. Inhibiting glutamine uptake represents an attractive new strategy for treating acute myeloid leukemia. Blood 2013, 122(20): 3521–3532.

7. Takahashi H, Inoue J, Sakaguchi K, Takagi M, Mizutani S, Inazawa J. Autophagy is required for cell survival under L-asparaginase-induced metabolic stress in acute lymphoblastic leukemia cells. Oncogene 2017, 36(30): 4267–4276.

8. Song P, Wang Z, Zhang X, Fan J, Li Y, Chen Q, et al. The role of autophagy in asparaginase-induced immune suppression of macrophages. Cell Death Dis 2017, 8(3): e2721.

9. Williams JC, Kizaki H, Weber G, Morris HP. Increased CTP synthetase activity in cancer cells. Nature 1978, 271(5640): 71–73.

10. Lynch EM, Hicks DR, Shepherd M, Endrizzi JA, Maker A, Hansen JM, et al. Human CTP synthase filament structure reveals the active enzyme conformation. Nat Struct Mol Biol 2017, 24(6): 507–514.

11. Lynch EM, Kollman JM. Coupled structural transitions enable highly cooperative regulation of human CTPS2 filaments. Nat Struct Mol Biol 2020, 27(1): 42–48.

12. Lin WC, Chakraborty A, Huang SC, Wang PY, Hsieh YJ, Chien KY, et al. Histidine-Dependent Protein Methylation Is Required for Compartmentalization of CTP Synthase. Cell Rep 2018, 24(10): 2733–2745 e2737.

13. Chakraborty A, Lin WC, Lin YT, Huang KJ, Wang PY, Chang IY, et al. SNAP29 mediates the assembly of histidine-induced CTP synthase filaments in proximity to the cytokeratin network. J Cell Sci 2020, 133(9).

14. Carra S, Seguin SJ, Lambert H, Landry J. HspB8 chaperone activity toward poly(Q)-containing proteins depends on its association with Bag3, a stimulator of macroautophagy. J Biol Chem 2008, 283(3): 1437–1444.

15. Ganassi M, Mateju D, Bigi I, Mediani L, Poser I, Lee HO, et al. A Surveillance Function of the HSPB8-BAG3-HSP70 Chaperone Complex Ensures Stress Granule Integrity and Dynamism. Mol Cell 2016, 63(5): 796–810.

16. Li F, Xiao H, Hu Z, Zhou F, Yang B. Exploring the multifaceted roles of heat shock protein B8 (HSPB8) in diseases. Eur J Cell Biol 2018, 97(3): 216–229.

17. Carra S, Seguin SJ, Lambert H, Landry J. HspB8 chaperone activity toward poly(Q)-containing proteins depends on its association with Bag3, a stimulator of macroautophagy. J Biol Chem 2008, 283(3): 1437–1444.

18. Arndt V, Dick N, Tawo R, Dreiseidler M, Wenzel D, Hesse M, et al. Chaperone-assisted selective autophagy is essential for muscle maintenance. Curr Biol 2010, 20(2): 143–148.

19. Cristofani R, Rusmini P, Galbiati M, Cicardi ME, Ferrari V, Tedesco B, et al. The Regulation of the Small Heat Shock Protein B8 in Misfolding Protein Diseases Causing Motoneuronal and Muscle Cell Death. Front Neurosci 2019, 13: 796.

20. Chang CC, Jeng YM, Peng M, Keppeke GD, Sung LY, Liu JL. CTP synthase forms the cytoophidium in human hepatocellular carcinoma. Exp Cell Res 2017, 361(2): 292–299.

21. Commisso C, Davidson SM, Soydaner-Azeloglu RG, Parker SJ, Kamphorst JJ, Hackett S, et al. Macropinocytosis of protein is an amino acid supply route in Ras-transformed cells. Nature 2013, 497(7451): 633–637.

22. Zhang J, Fan J, Venneti S, Cross JR, Takagi T, Bhinder B, et al. Asparagine plays a critical role in regulating cellular adaptation to glutamine depletion. Mol Cell 2014, 56(2): 205–218.

23. Lam SS, Martell JD, Kamer KJ, Deerinck TJ, Ellisman MH, Mootha VK, et al. Directed evolution of APEX2 for electron microscopy and proximity labeling. Nat Methods 2015, 12(1): 51–54.

24. Calise SJ, Carcamo WC, Krueger C, Yin JD, Purich DL, Chan EK. Glutamine deprivation initiates reversible assembly of mammalian rods and rings. Cell Mol Life Sci 2014, 71(15): 2963–2973.

25. Chang CC, Keppeke GD, Sung LY, Liu JL. Interfilament interaction between IMPDH and CTPS cytoophidia. FEBS J 2018, 285(20): 3753–3768.

26. Ji Y, Gu J, Makhov AM, Griffith JD, Mitchell BS. Regulation of the interaction of inosine monophosphate dehydrogenase with mycophenolic Acid by GTP. J Biol Chem 2006, 281(1): 206–212.

27. An S, Kumar R, Sheets ED, Benkovic SJ. Reversible compartmentalization of de novo purine biosynthetic complexes in living cells. Science 2008, 320(5872): 103–106.

28. Nakamura A, Nambu T, Ebara S, Hasegawa Y, Toyoshima K, Tsuchiya Y, et al. Inhibition of GCN2 sensitizes ASNS-low cancer cells to asparaginase by disrupting the amino acid response. Proc Natl Acad Sci U S A 2018, 115(33): E7776–E7785.

29. Wu CS, Li HP, Hsieh CH, Lin YT, Yi-Feng Chang I, Chung AK, et al. Integrated multi-omics analyses of oral squamous cell carcinoma reveal precision patient stratification and personalized treatment strategies. Cancer Lett 2025, 614: 217482.

30. Aviner R. The science of puromycin: From studies of ribosome function to applications in biotechnology. Comput Struct Biotechnol J 2020, 18: 1074–1083.

31. Andrejeva G, Gowan S, Lin G, Wong Te Fong AL, Shamsaei E, Parkes HG, et al. De novo phosphatidylcholine synthesis is required for autophagosome membrane formation and maintenance during autophagy. Autophagy 2020, 16(6): 1044–1060.

32. Ke PY, Chang CW, Hsiao YC. Baicalein Activates Parkin-Dependent Mitophagy through NDP52 and OPTN. Cells 2022, 11(7).

33. Rusmini P, Cristofani R, Galbiati M, Cicardi ME, Meroni M, Ferrari V, et al. The Role of the Heat Shock Protein B8 (HSPB8) in Motoneuron Diseases. Front Mol Neurosci 2017, 10: 176.

34. Wang PY, Lin WC, Tsai YC, Cheng ML, Lin YH, Tseng SH, et al. Regulation of CTP Synthase Filament Formation During DNA Endoreplication in Drosophila. Genetics 2015, 201(4): 1511–1523.

35. Pai LM, Wang PY, Lin WC, Chakraborty A, Yeh CT, Lin YH. Ubiquitination and filamentous structure of cytidine triphosphate synthase. Fly (Austin) 2016, 10(3): 108–114.

36. Liu S, Yao S, Yang H, Liu S, Wang Y. Autophagy: Regulator of cell death. Cell Death Dis 2023, 14(10): 648.

37. Iba T, Helms J, Maier CL, Ferrer R, Levy JH. Autophagy and autophagic cell death in sepsis: friend or foe? J Intensive Care 2024, 12(1): 41.

38. Chang HC, Tsai CY, Hsu CL, Tai TS, Cheng ML, Chuang YM, et al. Asparagine deprivation enhances T cell antitumour response in patients via ROS-mediated metabolic and signal adaptations. Nat Metab 2025, 7(5): 918–927.

39. Perez-Riverol Y, Bai J, Bandla C, Garcia-Seisdedos D, Hewapathirana S, Kamatchinathan S, et al. The PRIDE database resources in 2022: a hub for mass spectrometry-based proteomics evidences. Nucleic Acids Res 2022, 50(D1): D543–D552.

