## Supplementary figures for "HSPB8 regulates CTP synthase filaments to couple nucleotide metabolism and autophagy in tumors"

Chun-Yen Wang *et al.*

**This PDF file includes:**

Figs. S1 to S8

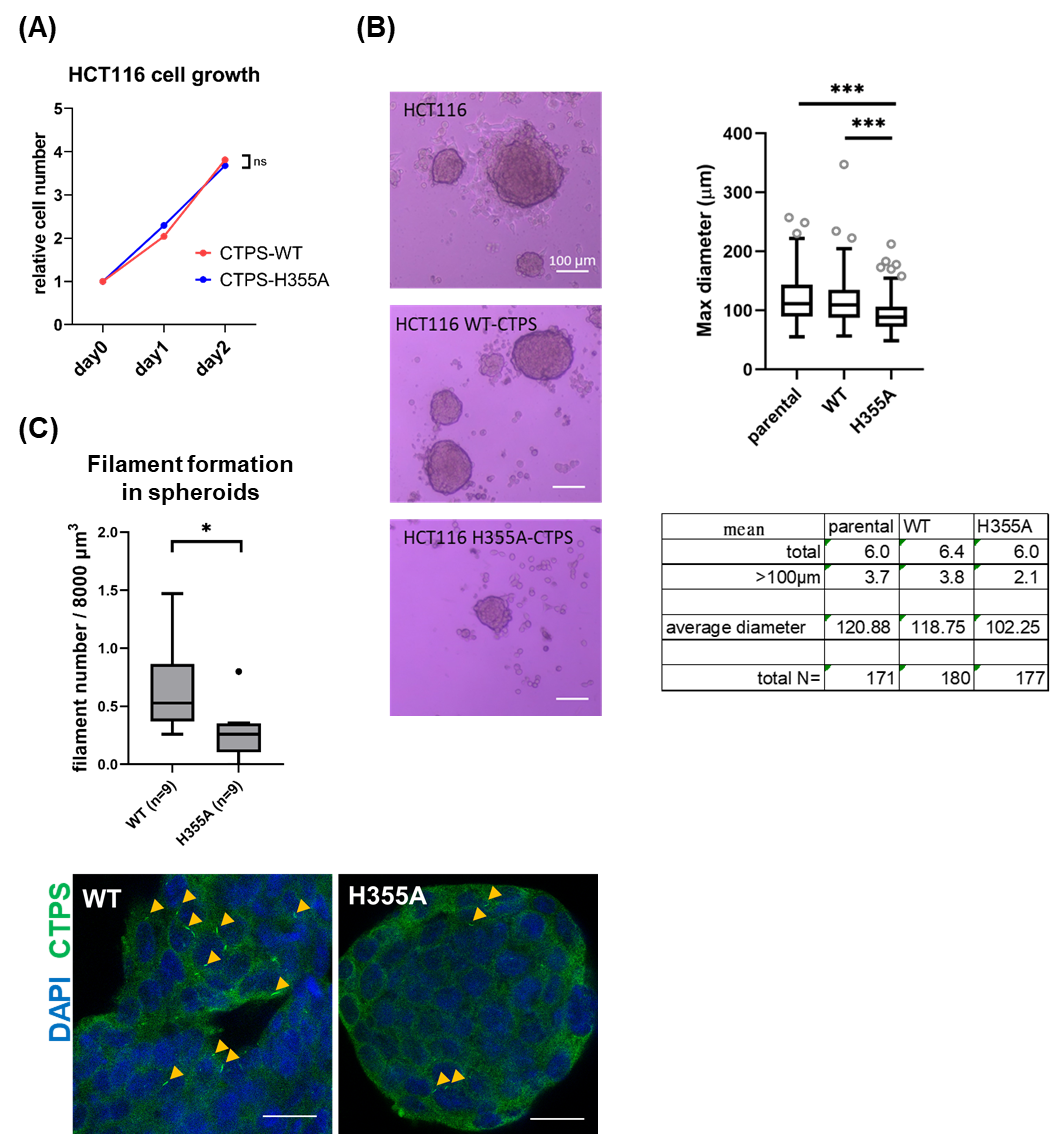
**Supplementary Figures**

**Supplementary Figure 1. CTPS filamentation provides growth benefits in 3D spheroid but not in 2D monolayer proliferation**

**(A)** Cell growth under standard culture conditions in McCoy’s 5A medium. **(B)** Representative images of spheres cultured under identical conditions for 7days (left). Spheroid diameter distribution is quantified in the right panel, with detailed values summarized in the table below. Total: the mean of total number of spheroids per 9.6 cm^2^ well; >100μm: the mean number of spheroids with diameter greater than 100μm per 9.6 cm^2^ well. Total N: the total number of spheroids counted. **(C)** Representative confocal images of HCT116 spheroids expressing wild-type CTPS (CTPS^WT^) or filament-null mutant (CTPS^H355A^) are shown in the bottom panel. Sections were immunostained for CTPS (green) and counterstained with DAPI (blue); yellow arrowheads indicate representative CTPS filaments. The top chart quantifies the mean filament density per 8000^3^. Scale bars: 20 mm.

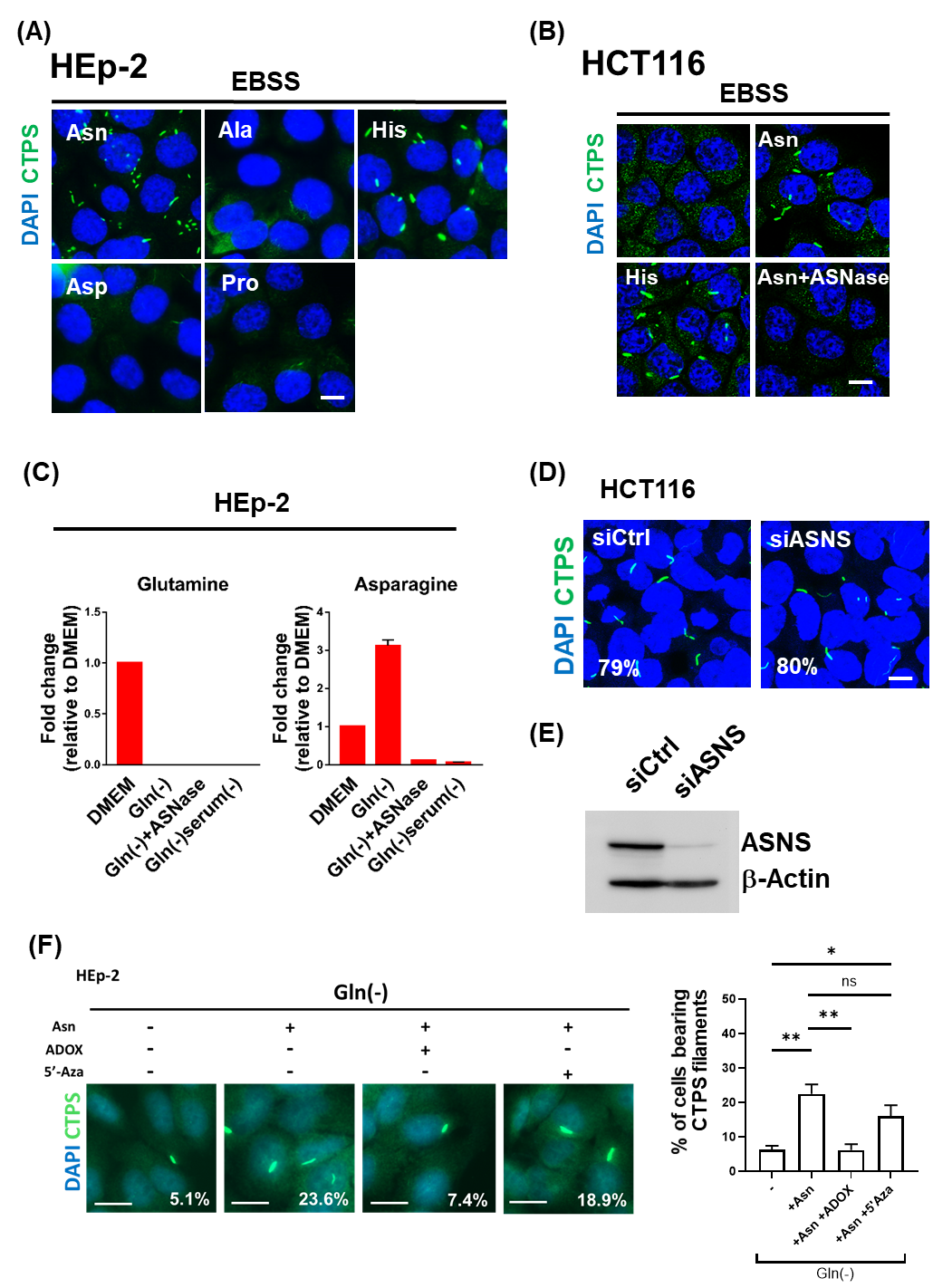
**Supplementary Figure 2. Regulation of CTPS filaments by serum-sourced asparagine and cellular protein methylation**

**(A)** HEp-2 cells were incubated with Earle’s balanced salt solution (EBSS) supplemented with histidine (His), alanine (Ala), asparagine (Asn), aspartate (Asp), or proline (Pro) for 6 h, followed by immunostaining for CTPS (green);nuclei were counterstained with DAPI (blue). **(B)** HCT116 cells were incubated in EBSS, supplemented with His, Asn, or asparaginase (ASNase) plus Asn for 6 h, followed by immunostaining for CTPS (green) and DAPI (blue). **(C)** HEp-2 cells (8 × 10⁵) were seeded in DMEM and cultured overnight. The medium was then replaced with Gln-free DMEM, serum-and Gln-free medium or Gln-free medium supplemented with ASNase for 24 h. Intracellular Gln and Asn levels were quantified. **(D and E)** HCT116 cells were transfected with scramble control or ASNS siRNA for 48 h, and then incubated in Gln-free DMEM medium for 24 h. Cells were subjected to immunostaining for CTPS (green) and DAPI (blue), and harvested for Western blotting analysis. **(F)** HEp-2 cells were incubated in Gln-free DMEM supplemented with Asn (0.34 mM), Asn plus adenosine dialdehyde (Adox; 20 μM), or Asn plus 5-azacytidine (5’Aza; 20 μM) for 24 h, followed by immunostaining for CTPS (green) and DAPI (blue). Quantification shown in the right panel represents three biological replicates. P-value *<0.05, **<0.01, ***<0.001. Scale bars: 10 mm.

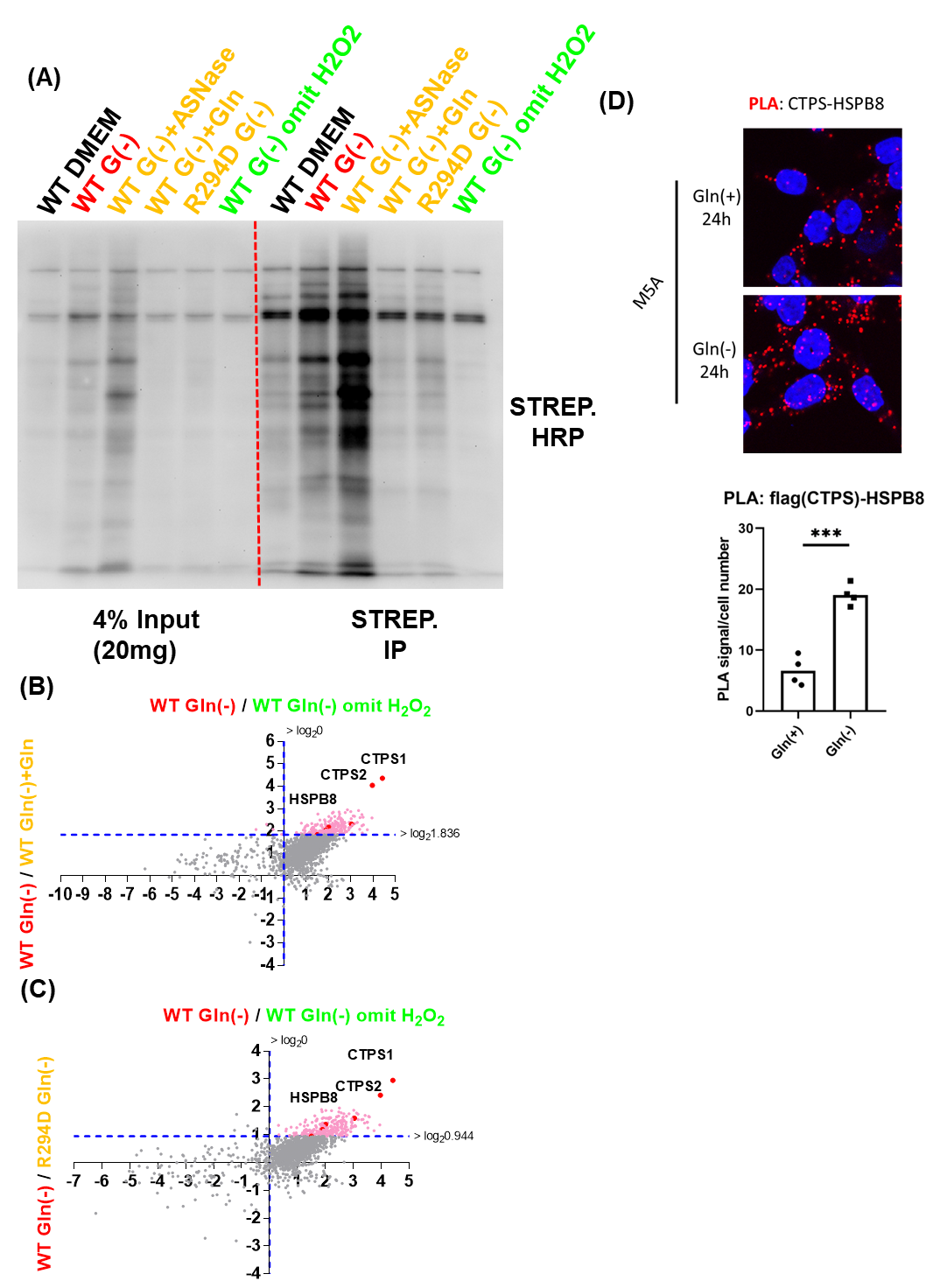
**Supplementary Figure 3. Validation of CTPS-APEX2 proximity labeling and verification of the CTPS–HSPB8 interaction**

**(A**) HEp-2 cells expressing CTPS^WT^ and CTPS^R294D^ were incubated under indicated conditions for 24 h. Biotinylated proteins were affinity purified using streptavidin-conjugated magnetic beads and detected by immunoblotting with streptavidin-HRP. **(B and C)** Scatter plots showing log2(WT G(-)/WT G(-)+Gln) versus log2(WT G(-)/WT G(-) without H_2_O_2_) and log2(WT G(-)/R294D G(-)) versus log2(WT G(-)/WT G(-) without H_2_O_2_), respectively, from CTPS filaments proximity labeling proteomic experiments. **(D)** Proximity ligation assay (PLA) between Flag-CTPS and HSPB8 in HCT116 cells cultured in control or Gln-free DMEM for 24 h.

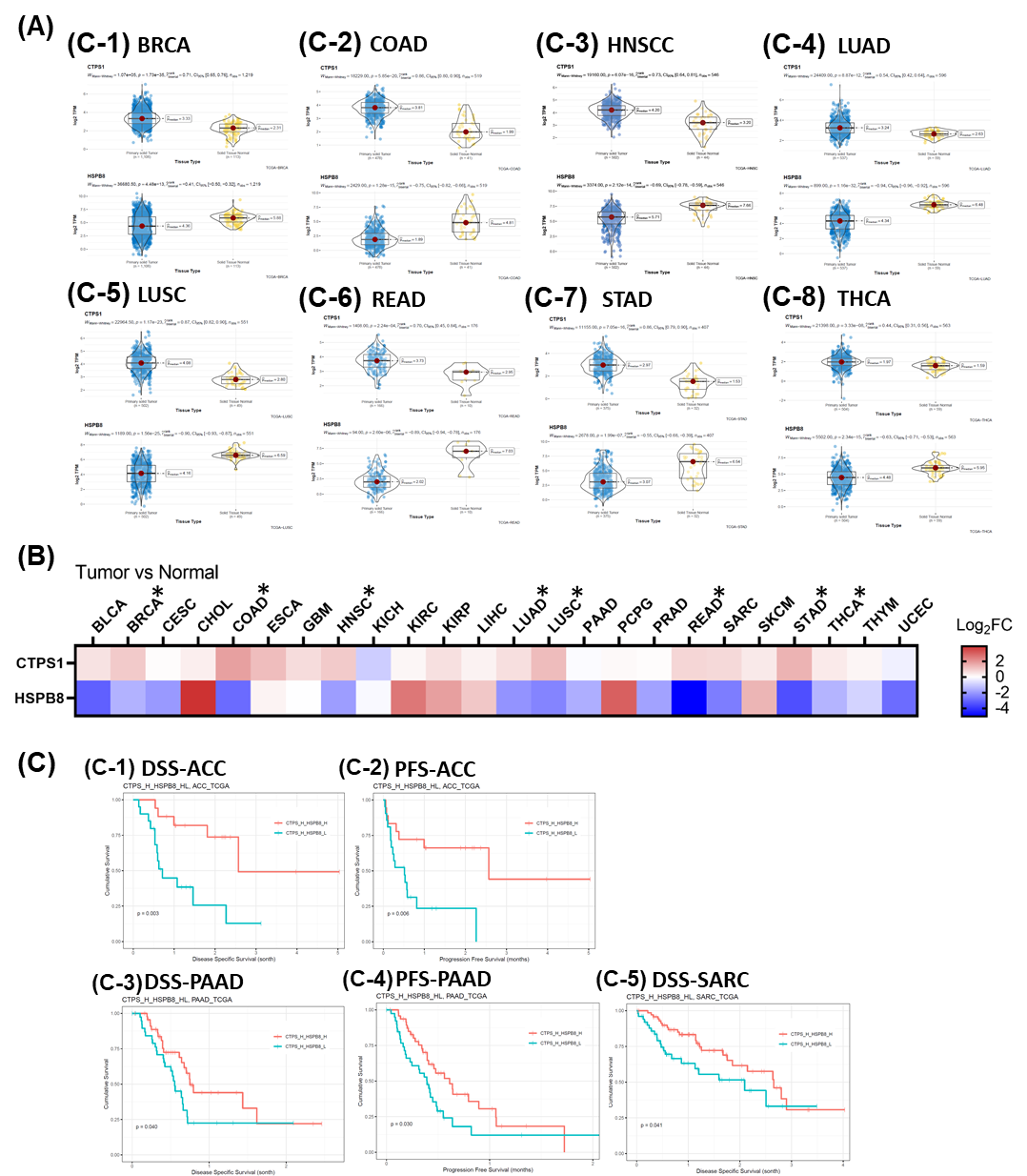
**Supplementary Figure 4. Anticorrelated CTPS and HSPB8 expression predicts poor survival in human cancers**

**(A)** Box plots show CTPS1 (top) and HSPB8 (bottom) expression levels (log₂ TPM) in tumor (blue) and normal (yellow) samples.

**(B)** Summary of data in (A). Analysis of TCGA tumor and matched normal tissue datasets showing an inverse relationship between CTPS and HSPB8 expression across eight cancer types (asterisks). Data are presented as log₂ fold change (tumor versus normal) of gene expression ratios.

**(C)** Kaplan–Meier disease-free (DSS) and progression-free (PFS) survival analysis indicating that tumors with high CTPS and low HSPB8 expression (cyan) are associated with significantly reduced survival, compared to that with high CTPS and high HSPB8 expression (red) in ACC (panel 1,2), PAAD (panel 3, 4), and SARC (panel 5).

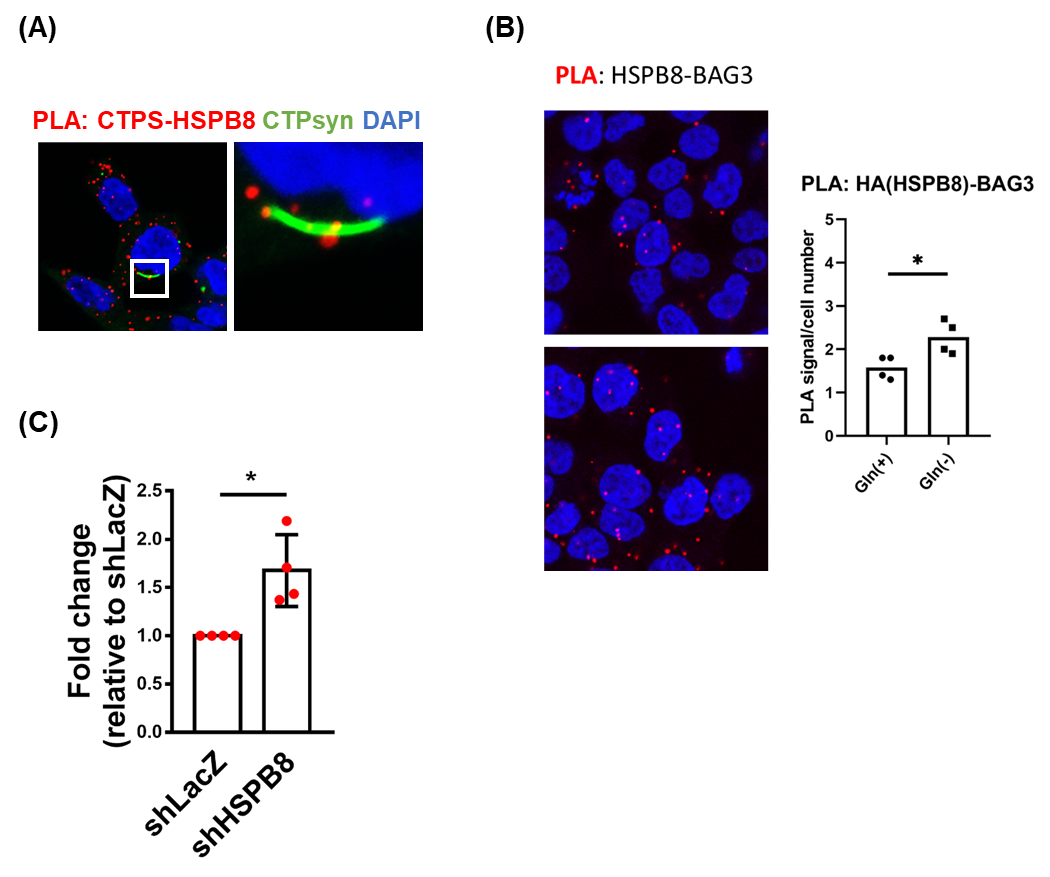
**Supplementary Figure 5. HSPB8 associates with CTPS filaments and BAG3 under glutamine deprivation**

**(A)** HCT116 cells stably expressing CTPS–GFP were cultured in Gln-free medium for 24 h, followed by proximity ligation assay (PLA) to detect interactions between CTPS and HA-HSPB8.

**(B)** HCT116 cells were cultured in Gln-free medium for 24 h, followed by PLA to assess interactions between HA-HSPB8 and BAG3.

**(C)** Quantification of puromycin incorporation shown in Fig. 5D, presented as fold change relative to control.

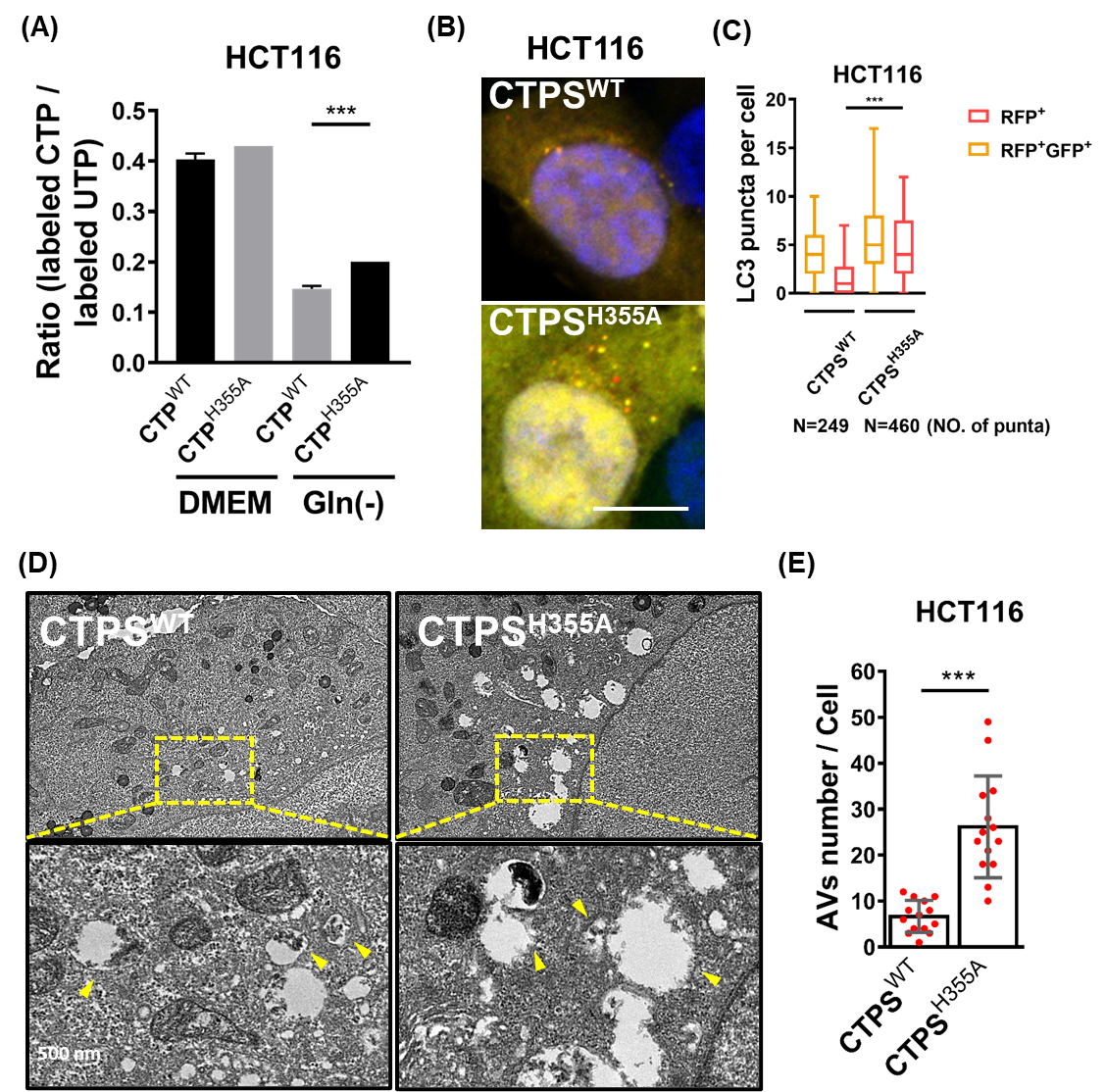
**Supplementary Figure 6. CTPS catalytic activity links nucleotide synthesis to autophagy under Glutamine deprivation**

**(A)** HCT116 cells stably expressing CTPS^WT^ and CTPS^H355A^ were cultured in DMEM or Gln-free media for 24 h, followed by treatment with ^13^C^15^N-uridine (100 mM) for 1 h. The ratio of labeled CTP to labeled UTP is shown. **(B and C)** HCT116 cells stably expressing CTPS^WT^ and CTPS^H355A^ together with the tandem GFP–RFP–LC3 reporter were cultured in Gln-free DMEM for 24 h. GFP- and RFP-positive puncta were quantified using ImageJ. **(D and E)** Transmission electron microscopy images of CTPS^WT^ and CTPS^H355A^ HCT116 cells cultured in Gln-free DMEM for 24 h. Autophagic vesicles are indicated by yellow arrowheads.

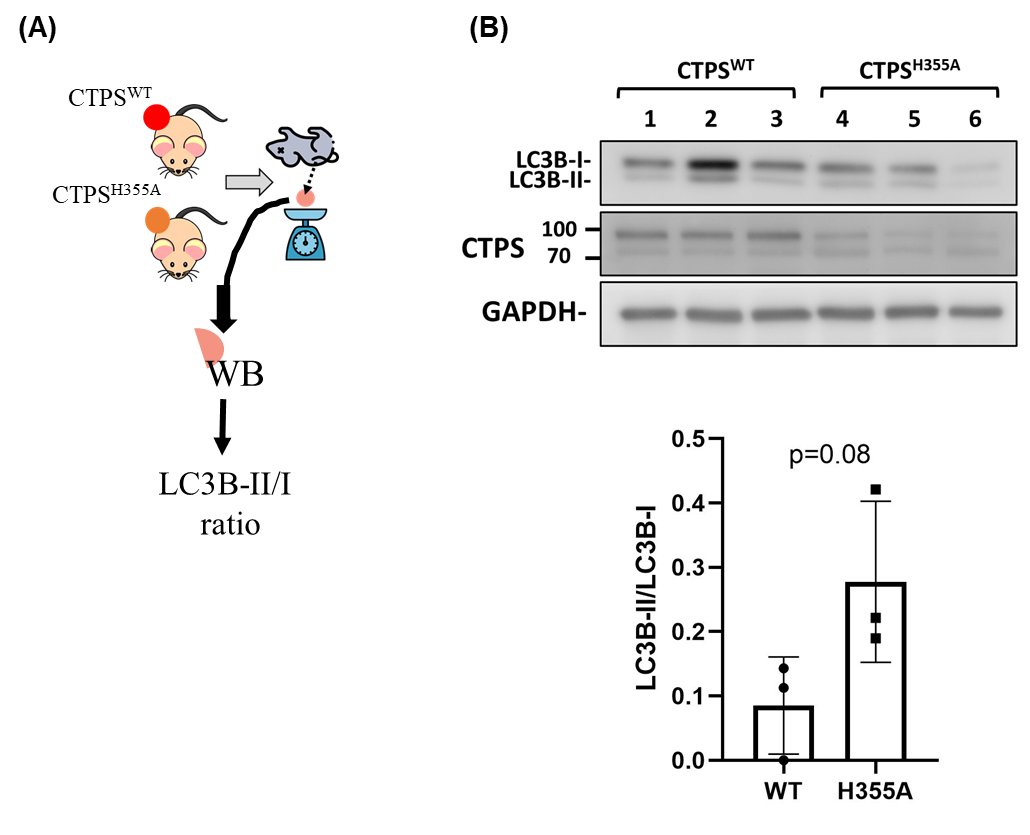
**Supplementary Figure 7. CTPS filament formation regulates autophagy in mouse tumors**

**(A)** CTPS^WT^ and CTPS^H355A^ mouse tumors (from Fig. 1, day 28) were dissected western blot (WB) analysis of LC3B.

**(B)** Western blot analysis of LC3B, CTPS, and GAPDH in tumor lysates. The LC3B-II/I ratio is summarized in the right panel (n = 3 tumors per group).

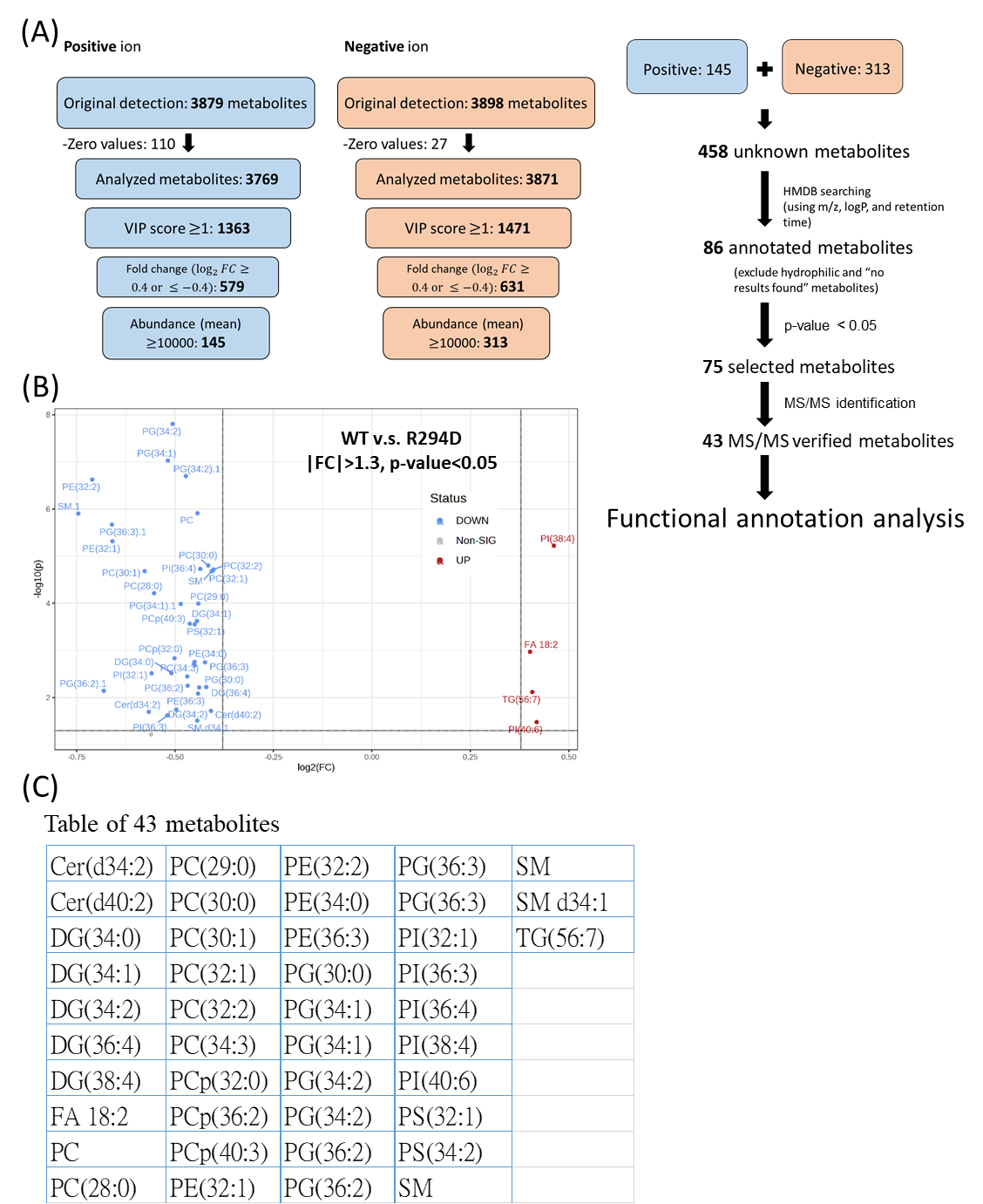
**Supplementary Figure 8. The strategy for lipid metabolite analysis**

**(A)** Flow chart of narrowing down strategy of lipid metabolites from LC-MS results. **(B)** Volcano plot of 43 identified metabolites. **(C)** Table of 43 identified metabolites.
